# Deconstructing the mouse olfactory percept through an olfactory ethological atlas

**DOI:** 10.1101/2020.11.09.374637

**Authors:** Diogo Manoel, Melanie Makhlouf, Charles J. Arayata, Abbirami Sathappan, Sahar Da’as, Doua Abdelrahman, Senthil Selvaraj, Reem Hasnah, Joel D. Mainland, Richard C. Gerkin, Luis R. Saraiva

**Author notes:** Correspondence and requests for materials should be addressed to R.C.G. or L.R.S.; twitter: @saraivalab).

## Abstract

Odor perception in non-humans is poorly understood. Here, we generated the most comprehensive murine olfactory ethological atlas to date, consisting of behavioral responses to a diverse panel of 73 odorants, including 12 at multiple concentrations. These data revealed that the mouse behavior is incredibly diverse, and changes in response to odor identity and intensity. Using only behavioral responses, ~30% of the 73 odorants could be identified with high accuracy (>96%) by a trained classifier. Mouse behavior occupied a low-dimensional space, consistent with analyses of human olfactory perception. While mouse olfactory behavior is difficult to predict from the corresponding human olfactory percept, three fundamental properties are shared: odor valence is the primary axis of olfactory perception; the physicochemical properties of odorants can predict the olfactory percept; and odorant concentration quantitatively and qualitatively impacts olfactory perception. These results provide a template for future comparative studies of olfactory percepts among species.

## INTRODUCTION

How sensory cues are translated into perceptual objects or complex behaviors remains a major unanswered question in neuroscience. Odor transduction in the nose leads to odor perception and to changes in behavior or physiology that are key for the species’ survival and reproduction, making the olfactory system an attractive model to address this question (1, 2).

In the last three decades, multiple studies have used the mouse as a model to elucidate basic molecular, cellular, and neural processes underlying mammalian olfaction (1). The increasing number of publicly available annotated genomes and recent advances in high-throughput sequencing technologies facilitated expanding this knowledge to human and other mammalian species, yielding new clues into the functional logic and the evolutionary dynamics of mammalian olfaction (3–5). On the other hand, most of our current knowledge about olfactory perception derives from human studies, facilitated by the generation and combination of large psychophysical datasets with chemoinformatic, statistical, and machine learning tools (6–11). These studies have yielded three key findings regarding the human olfactory percept. First, a large percentage of human olfactory perception variance is explained by a single dimension – odor valence (6, 7, 12). Second, the human olfactory percept for virtually any odorous molecule can now be predicted from chemical structure with decent accuracy (8). Third, odorant concentration can qualitatively alter perceived odor intensity and/or olfactory character (9, 13). But, do these findings also apply to the olfactory percept of other animals?

Characterizing olfactory perception in an animal relies on accurately quantifying multiple behaviors in response to large numbers of odorants, ideally at various concentrations. To date, most studies targeting olfactory perception in mouse include the scoring of only a single behavioral parameter (typically olfactory investigation time) upon exposure to a limited set of odorants (up to ~20), often delivered at different concentrations (14–18). A smaller number of studies have gone a step further by either scoring 10 behavioral parameters in response to a single odorant (19), up to 5 behavioral parameters upon exposure to 1-5 odorants (20–23), or by combining 3D-imaging with unsupervised machine learning techniques to identify specific sequences of mouse behavior in response to few odorants (24). We recently quantified the olfactory investigation time for 73 odorants delivered at 85 mM, and for a subset of 12 odorants at two additional descending concentrations (25). Despite these efforts, a systematic characterization of various mouse behaviors in response to a large panel of diverse odorants and several concentrations is still lacking. This prevents a systematic understanding of mouse olfactory behavior and how it relates to perception in humans and other species and limits our ability to study the neural computations underlying the transformation of odor stimuli at the nose to odor objects in the brain.

Here, we generated and investigated an atlas of odor-guided behaviors in mouse in response to a diverse panel of monomolecular odorants at different concentrations to deconstruct the mouse olfactory percept and unravel similarities and differences between olfactory perception in mouse and human.

## RESULTS

### The olfactory-ethological atlas

We recently generated a mouse behavioral video library and quantified the cumulative duration of olfactory investigation over a three-minute-long assay (3’dOI) to a panel of 73 odorants at 85 mM and an odorless control (i.e., water or H2O) (25). We classified the odorants as eliciting avoidance or approach if the 3’dOI was significantly lower or higher than H2O, respectively (25). Here, we exploited this video library by scoring 17 additional behavioral parameters (indicative of valence, stress, or exploration) from 410 individual mice exposed to 73 odorants and H2O (see Methods for details, Fig. 1A-D). Each mouse was exposed to only a single odorant, and thus the data for each odorant consisted of 5-10 mice, each scored on 18 behaviors.

**Figure 1.**
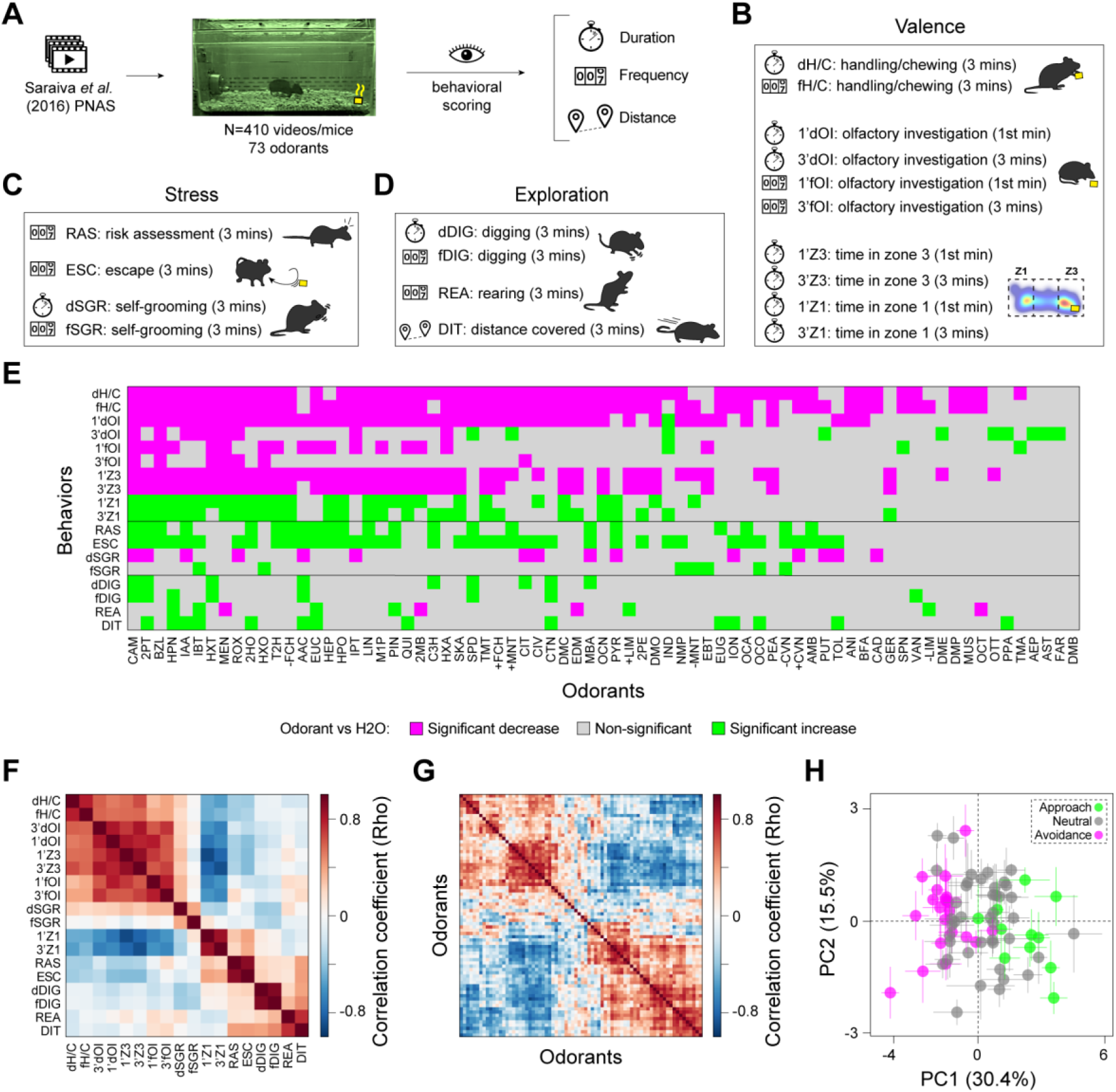
An olfactory-ethological atlas and the primary axis of olfactory perception in mouse. (**A**) Study design: videos were retrieved from (25) and scored for a total of 18 distinct behavioral parameters, which we grouped in three broad categories: (**B**) valence, (**C**) stress, (**D**) exploration. (**E**) A graphical display in the form of a heatmap summarizing the combinatorial behavioral codes for the 73 odorants tested. Behaviors showing significant (one-way ANOVA, BKY multiple comparisons correction, n = 5-10 per odorant) increases, decreases and non-significant responses compared to H2O are indicated in green, magenta and grey squares, respectively. (**F**) Correlation across odorants between each of the 18 behaviors (n = 3-7 per odorant). (**G**) Correlation across behaviors between each of the 73 odorants and H2O (n = 3-7 per odorant). In both A and B, odorants and behaviors are ordered to illustrate clustering. Two major clusters of odorants stand out. (H) Principal Component Analysis of the 18 mouse behavioral parameters for H2O and the 73 tested odorants (n = 3-7 per odorant). Circles are colored to indicate avoidance (magenta), neutral (grey), or approach (green) odorants. The bars indicate the standard error.

This analysis yielded a dataset totaling 7606 individual data points (Fig. S1, Data File S1), corresponding to individual-odorant-behavior triads, from which we also calculated the across-individual average value for each of the 1314 odorant-behavior pairs. Of these pairs, 470 (or 35.8%) are significantly different (p<0.05, one-way ANOVA, BKY multiple comparisons correction) from the H2O control, with 293 corresponding to decreases and 177 to increases of the duration, frequency, or distance of the behavioral measure (Fig. 1E, Data File S1). Of the 72 odorants eliciting at least one significant behavioral change, the vast majority (57 or 78%) exhibited unique combinations of significant increases and decreases among the 18 different behaviors, hereafter referred to as combinatorial behavioral codes. Together, these results indicate that combinatorial behavioral codes are incredibly diverse – with most odorants studied here eliciting a unique behavioral code

### Valence is the primary axis of olfactory perception in mouse

Pleasantness, a surrogate measure for odor valence, has emerged across multiple studies as the primary axis of human olfactory perception (7, 12, 26). However, the extent to which this is conserved in mouse remains unknown. We examined the structure of the behavioral response matrix (behaviors x odorants) to address this question. From this response matrix, we computed two correlation matrices (between behaviors, Fig 1F, and between odorants, Fig 1G) and arranged the ordering of each to match the results of hierarchical clustering. In both cases, there was clear evidence for two self-similar groups (upper left and lower right patches in Fig. 1F and Fig. 1G). Indeed, principal components analysis (PCA) showed that a single dimension can explain ~1/3 of the variance in the data (Fig. 1H and Fig. S2). The first dimension of PCA analysis cleanly separates the approached from the avoided odorants, which we interpret as a valence axis (Fig. 1H).

### Discriminability of odorants using behaviors

Odorants lie in stereotypical locations in a behavioral space defined by the first two principal components of the response matrix (Fig. 1H). We next asked how well odorants can be distinguished from one another using the original 18-dimensional space. We computed D’, a general measure of discriminability between two signals, for all pairs of odorants. Higher values of D’ indicate greater discriminability between two odorants using behavioral signals, with a value of 1.0 indicating that the magnitude of the difference in two such signals is equal to their variability across individual mice; it is thus also a measure of effect size.

We first used single behaviors as the signal with which to compute D’. We also computed D’ for shuffled data in which odorant labels were permuted across individual mice, but the same number of mice were “assigned” to each odorant. Each behavior had a mean D’ value (across odorant pairs) of between 0.6 and 1.0; by contrast, shuffling resulted in lower D’ values (0.5-0.6) (Fig. 2A), and every single behavior exhibited significantly higher D’ (over all odorants pairs) in the original than in the shuffled data (Fig. 2B).

**Figure 2.**
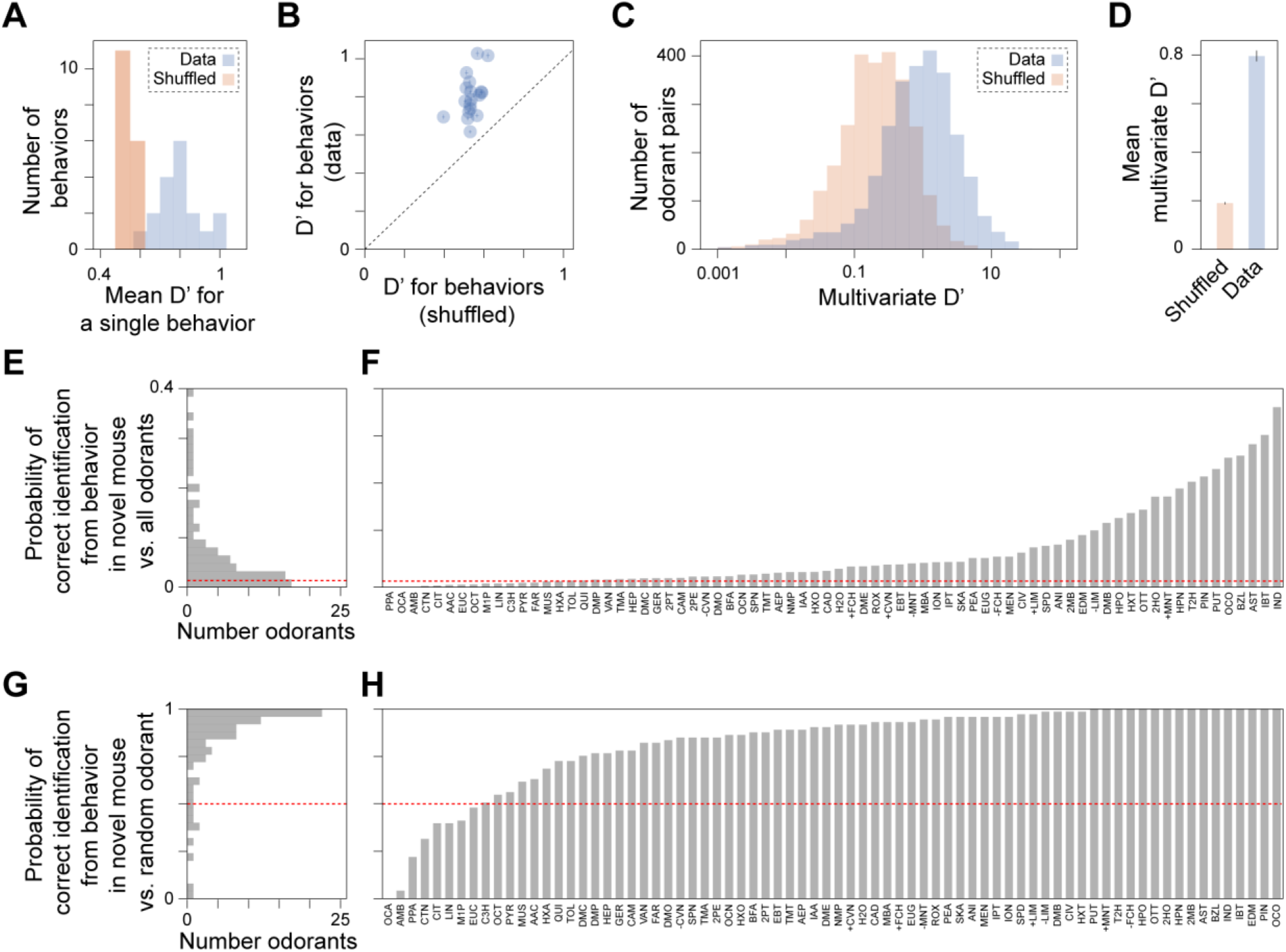
Discriminability of odorants using behaviors and out-of-sample prediction of odor identity. (**A**) D’, a measure of discriminability between two odorants, is greater for the data than for a shuffling (across mice) of the the data. (**B**) Same data as in A, except shown for each behavior (circle) vs its corresponding shuffle. Error bars (inside circles) represent SEM taken over all odorant pairs. (**C**) Using all behaviors simulantaneously, the multivariate measure D’ is computed. (**D**) Multivariate D’ is ~8x larger for the real data than for shuffled data. (E-H) A linear discriminant analysis classifier was trained on all odorants, using all but one mouse for each odorant. Predictive performance was evaluated for the remaining mice (one odorant each). (**E**) A histogram of the probability that the correct odorant (out of 74 possibilities) is identified from a new mouse’s behavior. The dashed red line reflects chance performance. (**F**) Mean performance for each odorant; higher values mean the odorant is easier to uniquely identify from behavior. (**G**) Similar to A, but for classification of the true odorant against a random alternative odorant. Chance is now 50%, as reflected by the dashed red line. (**H**) Mean performance for each odorant. 1.0 means that behavior was always sufficient to identify the odorant vs. any specific alternative odorant.

Because some behaviors may carry distinct and complementary information about odorant identity, we also computed a multivariate equivalent of D’, which answers the same question about odorant discriminability but using all behaviors simultaneously. Additionally, because this yields a single number for each odorant pair, we can compute the variability of this number across odorant pairs, which are presumably heterogeneous in their behavioral discriminability. This multivariate D’ is indeed heterogeneous across odorant pairs (Fig. 2C) but is on average 4x larger (8x larger in log-transformed space) for the real data than for shuffled data (Fig. 2D). If most of the behavioral variability was mouse-specific and not odor-specific, D’ would have been <<1. Thus, our observation of D’ ~1 indicates that this behavioral ensemble represents a general odor-evoked behavioral code in these mice.

While D’ is a common measure of discriminability, it overestimates discriminability when the number of replicates (individuals) per stimulus is small; this is why D’ for shuffled data is > 0. Instead, we can ask how accurately a trained statistical model can select the correct odorant from an out-of-sample observation of mouse behavior, i.e., a mouse on whose data it has not been trained. We trained a simple linear discriminant analysis classifier on all odorants, but withheld one mouse for each odorant to use for testing the performance of the classifier. We asked the classifier to perform two tasks: first, to predict the correct odorant (out of 74 possibilities) given a novel observation of behavior, and second, to predict the correct odorant (out of two choices: the correct odorant and one other chosen at random). In the first task, most odorants could be predicted at above chance (1/74) levels (Fig. 2E,F). Some could even be predicted correctly 30-40% of the time (IND, IBT, AST). In the second task, 66/74 odorants could be identified from behavior at above chance (½) levels against a random odorant, and 22/74 could be identified >96% of the time in the same comparison (Fig. 2G,H). Consequently, odor-evoked behavior for most odorants was stereotypical enough across individual mice to identify those same odorants from the behavior of novel mice.

### Reconstructing a low-dimensional space of mouse olfactory behavior

While we recorded 18 distinct behaviors and used them all in our analyses of discriminability and odorant prediction, many of these behaviors are correlated (Fig. 1F). Thus, the underlying dimensionality of the olfactory behavioral space may be much lower than 18. PCA indicated that 90% of the variance was explained by ten dimensions (Fig. S2), but this is likely to be an overestimate for two reasons: first, at least some of this variance is noise, driven by within-odorant variability in behavior, and cannot be removed by averaging; second, it is well-known that PCA does not produce the most natural decomposition of the data in many applications (27). To overcome the first concern, we asked how many dimensions are required to optimally represent each odorant’s behavioral phenotype. To address the second we used non-negative matrix factorization (NMF) (27), a decomposition technique known for producing compact, intuitive, representations of data in numerous domains, including olfactory perception (28). Specifically, we computed an NMF decomposition of the behavioral dataset and asked for what dimensionality of this decomposition maximized the intraclass correlation coefficient (ICC) using novel mice. The ICC measures the fraction of total behavioral variance (across all mouse-odorant pairs) that is explained by odorant identity. Theory suggests that a low-dimensional NMF decomposition might effectively denoise behavioral data by identifying and discarding noisy, irrelevant dimensions, thus increasing ICC. However, if the decomposition compresses too many dimensions, odorant-associated structure could be lost, reducing the ICC. Thus, the dimensionality at which the ICC is largest reflects the most natural dimension for preserving information about olfactory behavior.

We found that ICC was greatest for a 2-dimensional space (ICC=0.57, Fig. 3A), substantially higher than for the original 18-dimensional space (ICC=0.34) or a 1-dimensional space (ICC=0.35). In other words, a 2-dimensional space was optimal for explaining behavioral variance across experiments in terms of the odorants used as stimuli. In contrast, shuffling odorant labels across experiments produced a consistently low value (ICC=0.2) across all values of dimensionality. To understand the meaning of these two dimensions, we examined the weights in the NMF decomposition for each behavior in each of these dimensions (Fig. 3B). We observe that the behaviors with the highest weight correspond to the valence-related category (handling/catching – H/C, olfactory investigation – OI and zone assays – Z1/Z3), followed by the exploratory (rearing – REA, digging – DIG, distance covered – DIT) and stress related categories (risk assessment – RAS, and escape – ESC). These results are consistent with what was observed using PCA, but goes further by providing a parts-based understanding of the fundamental units of olfactory behavior in mice.

**Figure 3.**
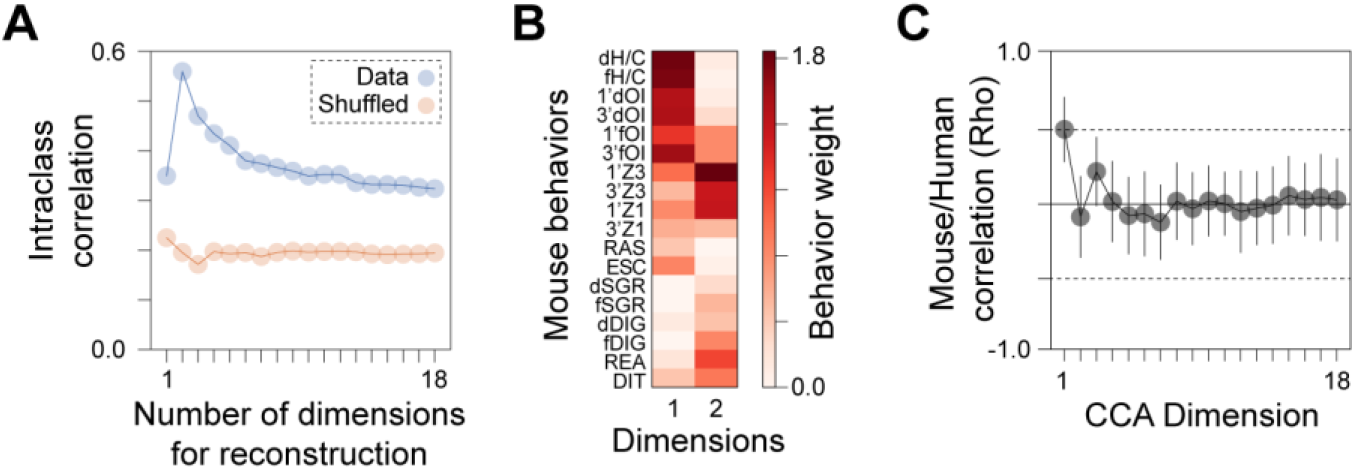
Reconstructing mouse behavior and the alignment between mouse and human behavioral spaces. (**A-B**) The optimal reconstruction of mouse behavior requires few dimensions. (**A**) Non-negative matrix factorization (NMF) is used to learn a low-dimensional (≤ the number of measured behaviors) representation of the behavioral data. The intra-class correlation coefficient (ICC), reflecting the behavioral agreement within (vs. across) odorants is shown, as a function of the number of dimensions used. Lower numbers of dimensions effectively denoise the data. Results for the data are shown in blue, results for the data with shuffled odorant labels are shown in orange. Eighteen dimensions would reflect independent contributions of each behavior. ICC is maximized for a 2-dimensional representation of behavior. (**B**) Contributions of behaviors to the resulting 2 dimensions. (C-D) Alignment of mouse and human behavioral spaces. (**C**) Canonical correlation analysis co-aligns mouse behavioral features and human-provided descriptors for the same odorants. Canonical dimensions were computed using all but one odorant, and the remaining odorant was used to evaluate the correlation (Pearson) between mouse behavior and human percepts. Error bars represent standard deviation across held-out odorants. P-values were computed by comparing to shuffled data.

### Mouse vs. Human

Comparative studies of odor valence in mice and humans are scarce and have yielded conflicting results (17, 29). Indeed, whether mouse and human share the same olfactory preferences, and more importantly, the same olfactory percept, is still unknown. To address this gap, we first compared the mouse behavioral dataset to the human-rated intensity and pleasantness reported in a previous study (9). For 23 overlapping odorants between both studies, we found no significant correlation between any of the 18 mouse behavioral parameters and human-rated intensity or pleasantness (Fig. S3), consistent with our previous study (29). We then extended this analysis to include 19 additional human semantic descriptors (e.g., fish, sweet, chemical) for (9), but found only seven significant correlations (out of a possible 475) encompassing five mouse behaviors and four human semantic descriptors.

To identify the shared structure between mouse olfactory behavior and human olfactory perception, 23 odorants may have been insufficient. To make use of a larger number of odorants, we used the data and machine-learning algorithms from the DREAM Olfaction Prediction Challenge (8) to obtain predictions for the human-rated intensity, pleasantness, and 19 semantic descriptors for the 73 odorants tested in our mouse experiments (see Methods). Rather than identify simple correlations between mouse behaviors and human perceptual descriptors, we asked whether mouse behavioral space and human perceptual space could be projected onto a common basis. If so, they might simply be two views of a common mammalian olfactory perceptual space. We used canonical correlation analysis (CCA) to obtain this basis using all but one odorant, and then asked whether this basis could identify shared structure in the perceptual/behavioral space of these two species using the remaining (out-of-sample) odorant. Our results (Fig. 3C) identified a single dimension in which these two spaces were correlated (Pearson’s rho=0.50), with additional dimensions failing to capture any additional shared structure.

### Predicting behavior from chemical structure

Prior studies have identified relationships between certain molecular features of aliphatic odorants, such as hydrocarbon chain length or chemical functional group, and their induced neural responses or perceptual similarity (30–32). Of the twelve aliphatic odorants in our dataset we found no apparent association between chemical functional groups and valence (Fig. 4A) but observed a strong positive correlation between hydrocarbon chain length and 3’dOI (Spearman’s rho, rs = 0.795, P = 0.0034; Fig. 4B, Data File S2). Concomitantly, we observed similar significant relationships for three additional valence-related olfactory investigation parameters (1’dOI, 1’fOI, and 3’fOI), but not for any of the remaining 14 behavioral parameters. These results suggest that the hydrocarbon chain length of aliphatic odorants could be tied to their valence in mouse. While interesting, due to the limited number of associations tested, the above results are not generalizable. Indeed, the olfactory percept of an odorant is typically not explainable using only restricted feature metrics like the ones analyzed above, but instead, larger sets of physicochemical descriptors are needed (7, 33). In this context, robust correlations have been identified between large sets of physicochemical descriptors and multiple olfactory perceptual qualities in human, and also olfactory investigation time in mouse (8, 9, 17, 34).

**Figure 4.**
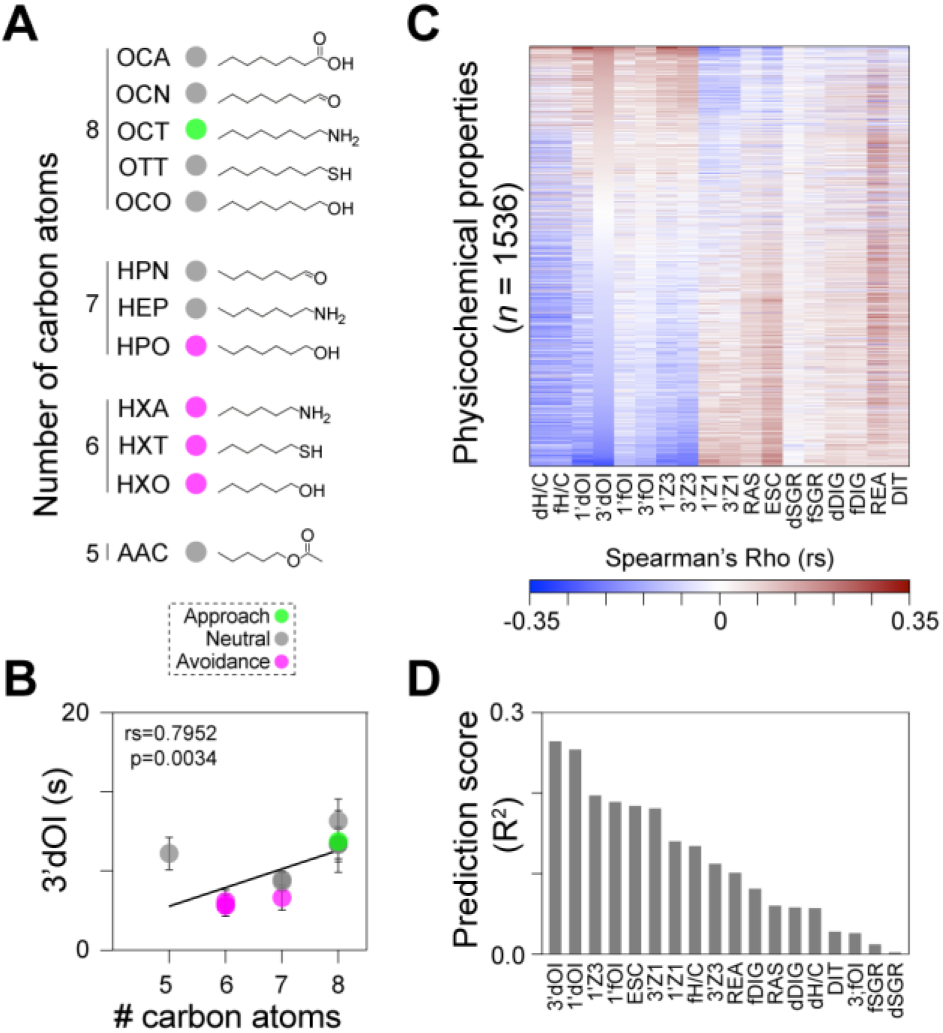
Predicting mouse olfactory-driven behaviors. (A) The twelve n-aliphatic odorants in our dataset include molecules with varying lengths (5-8 carbon atoms) and different functional groups (amino, thiol, hydroxyl, carboxyl, ester or aldehyde), amyl acetate (AAC), hexanethiol (HXT), hexylamine (HXA), hexanol (HXO), heptanol (HPO), heptanal (HPN), heptylamine (HEP), octanoic acid (OCA), octanol (OCO), octanal (OCN), octylamine (OCT), and octanethiol (OTT). Circles are colored to indicate aversive (magenta), neutral (grey), or approached (cyan) odorants. (B) Correlation plot between odorants with different numbers of carbon atoms and the total duration of olfactory investigation (3’dOI). Data points indicate mean, and error bars show SEM (n = 5-10 per odorant). (C) Heatmap depicting Spearman’s correlation coefficient (rs) between 1536 physicochemical descriptors and the 18 mouse behavioral parameters. The physicochemical descriptors are sorted in descending order of the rs for 3’dOI. (D) Performance of the support vector machine model on the 18 behavioral parameters.

To test whether different mouse behaviors correlate to distinct sets of physicochemical descriptors, we started by retrieving 4,870 physicochemical descriptors for each of the 73 odorants (plus water) using Dragon 6.0 (Talete) software. After removing descriptors with near-zero variance or missing data, we next calculated the Spearman’s correlation coefficient of the remaining subset of 1,536 physicochemical descriptors against the 17 behavioral parameters for all 410 individual mice and all odorants tested, including water (Fig. 4C and Data File S2). Of the possible 27,648 interactions, 29.5% (or 8,141) resulted in significant (p<0.05) interactions, with the duration of self-grooming (dSGR) and duration of handling/catching the stimulus (dH/C) being the parameters eliciting the minimum (16) and maximum (968) of significant interactions.

Multiple studies in humans have used specific subsets of physicochemical properties to reverse-engineer the pleasantness, intensity, and other odor qualities of odorants (8, 34, 35). One question raised by these studies and our results above is whether olfactory-driven behaviors in the mouse can also be predicted by the chemical structure of odorants. To answer this question, we used an unsupervised machine-learning approach (see Methods) to determine the amount of behavioral variance that could be explained by the physicochemical properties of the odorants. To further reduce redundancy among the 1,536 physicochemical descriptors (i.e., features) from the step above, we removed any descriptors sharing a correlation ≥95% with any other descriptor, which left us with a final total of 662 features to use in our downstream analysis. We found that the mean predictive accuracy (R2) for each behavior varied greatly, with dSGR displaying the lowest average test set prediction (R2=0.003) score, and 3’dOI displaying the highest average test set predictions (R2=0.265) score, respectively (Fig. 4D, Data File S2).

### Behavioral effects of odorant concentration

In human, the quality, valence, and intensity of odorants can change with the concentration (11, 13, 36). In mouse, the valence (25, 37) and intensity (38) of odorants can also be concentration-dependent, but whether different concentrations of a given odorant can affect other odor-guided mouse behaviors remains largely unexplored. To address this question, we again exploited the video library from our previous study (25) and scored 123 videos from mice exposed to a subset of 12 odorants at two additional descending concentrations, 100x (850 μM) and 10,000x (8.5 μM) diluted from the concentrations used in the previous experiments (Data File S3). At the highest concentration, odorants are diverse in their valence, with some eliciting avoidance (IBT, IAA, TMT), no change (PEA, 2HO, DMP, VAN, SKA, AMB), or approach (IND, TMA, PUT) when presented at the original 85 mM concentration (Fig. 1A). By contrast, all of these odorants are neutral or elicit approach when tested at lower concentrations (Fig. 5A) (25). The scoring of the 17 other behaviors for this dataset yielded a total of 3474 individual data points, of which 2159 were newly scored for the new 850 μM and 8.5 μM concentrations (Data File S3). Of the 12×3×18 = 648 odorant-concentration-behavior pairwise comparisons, 117 (or 18.1%) were significantly different than the water control, with 42 corresponding to decreases and 75 to increases (Fig. 5A, Fig. S5A, Data File S3). The behaviors eliciting the highest (23) and lowest (1) number of significant changes were 3’dOI and dSGR, respectively (Fig. 5A, Data File S3), consistent with odor valence driving the largest fraction of behavioral variation. Of the 12 odorants tested, 10 elicited significant behavioral changes for all concentrations tested, while the remaining 2 failed to trigger significant behavioral changes for one of the concentrations (850 μM for DMP and 8.5 μM for PEA). Of the 3 concentrations tested across all odorants, 85 mM elicited the highest number of combined significant behavioral changes (60), followed by 8.5 μM and 850 μM, with elicited a total of 30 and 27 changes in behavior, respectively. We also identified 28/36 unique behavioral combinations in this dataset: 12 for 85 mM, 8 for 850 μM, and 8 for 8.5 μM (Fig. 5A).

**Figure 5.**
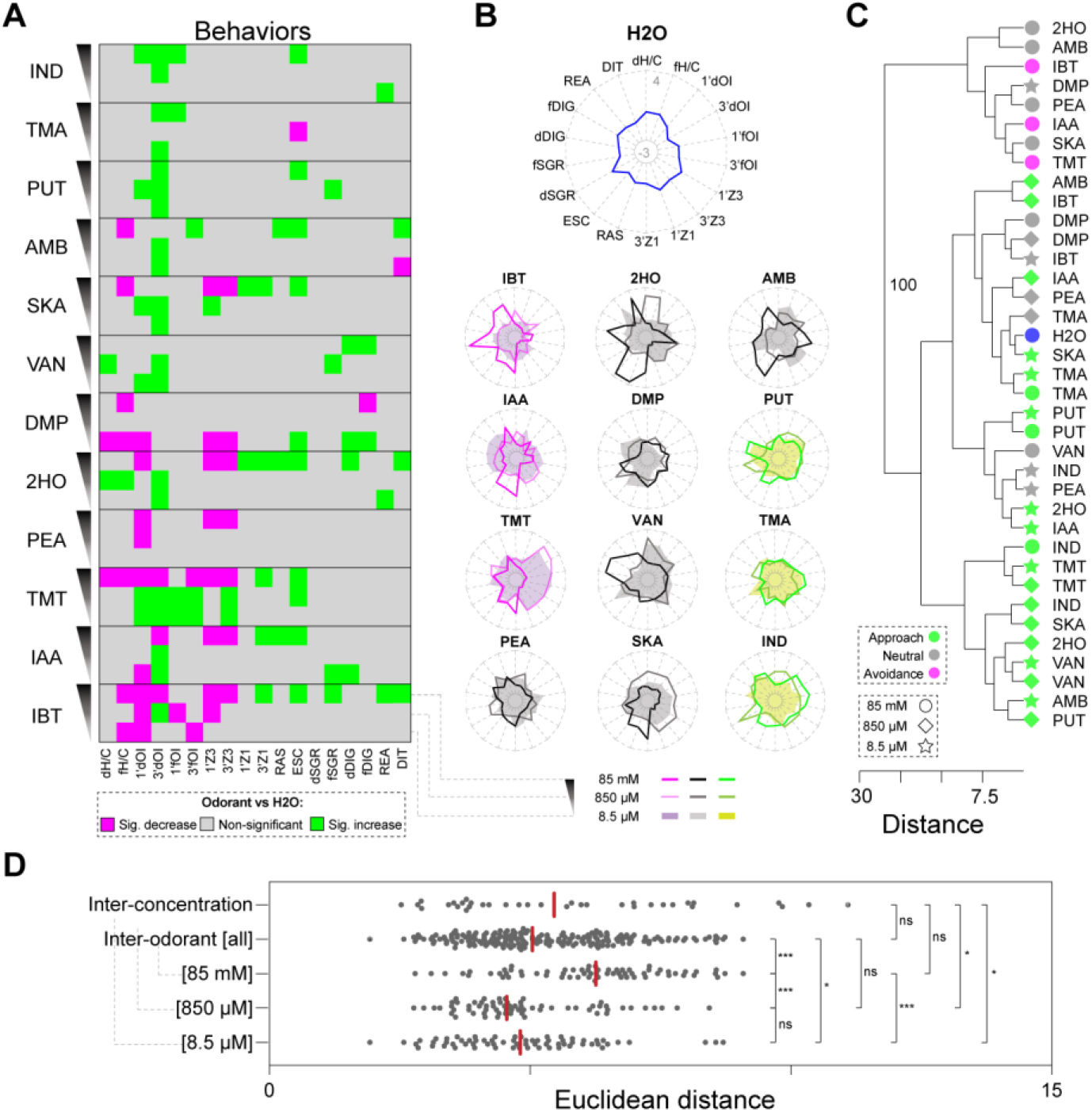
The effect of odorant concentration on behavior. (**A**) A graphical display in the form of a heatmap summarizing the behavioral codes for the 12 odorants tested at three different concentrations. Behaviors showing significant (one-way ANOVA, BKY multiple comparisons correction, *n* = 5-9 per odorant) increases, decreases, and non-significant responses compared to H2O are indicated in green, magenta and greye squares, respectively. Of the 12 odors tested, at 85 mM, three are aversive (TMT, IAA, IBT), six are neutral (PEA, SKA, VAN, DMP, 2HO, AMB), and the remaining three approached (IND, TMA, PUT,). (**B**) The behavioral profiles (radar plots, z-scores) for water (top center) and the 12 odorants tested at three concentrations (85 mM, 850 μM and 8.5 μM) are shown. The behavioral profiles for the odorants classified at 85 mM as aversive, neutral, and approached are shown in different shades of magenta, grey/black, and green, respectively. The corresponding descending concentrations are depicted in lighter hues of the same color, irrespective of their valence at 850 μM and 8.5 μM. (**C**) Hierarchical clustering analysis of the behavioral profiles for the 12 odorants tested at the three different concentrations supported the existence of 2 clusters. Bootstrap value (100 bootstraps) for the major node is indicated. (**D**) Plot comparing the Euclidean distances calculated for all pairwise inter-concentration comparisons (restricted to different concentrations within each odorant), inter-odorant comparisons (restricted to different odorants for each of same concentrations), and subsets of inter-odorant comparisons for each concentration separately (85mM, 850μM, and 8.5μM). Asterisks indicate significant differences (one-way ANOVA, BKY multiple comparisons correction): ns, non-significant; *P < 0.05; ***P < 0.001.

Next, we performed an ANOVA to estimate the role of concentrations as well as odorant-concentration interactions in generating behavioral responses. This analysis showed a main effect of either concentration or an interaction between odorant and concentration for nearly all behavioral parameters (p<0.005 for 16/18; p<10-8 for 13/18). The effect size for odorant identity (η2 = 0.27 +/− 0.05) was slightly greater than for odorant concentration (η2 = 0.18 +/− 0.04), but the interaction between odorant identity and concentration was stronger than either one alone (η2 = 0.43 +/− 0.04).

The results above showed that even at low concentrations, odorants can significantly impact mouse behavior and revealed a remarkable diversity in the behavioral responses among the different odorants and across concentrations. However, these results raise other important questions: How exactly does mouse olfactory behavior change with concentration? Is odor identity preserved across concentrations? And what has a more significant impact on the mouse olfactory percept: odorant identity or odorant concentration?

To answer these questions, we first performed a comparative analysis of the behavioral profiles for the different odorants and concentrations tested. We found that the highest concentration tested (85 mM) yields the most distinguishable behavioral profiles across all odorants, and noted similarities between the behavioral profiles of the same odorants at the two lower concentrations of 850 μM and 8.5 μM (Fig. 5B). This overlap was particularly noticeable between the behavioral profiles of those two lower concentrations for TMA, TMT, and VAN. We also observed other common trends among all odorants as we decreased their concentration: while the scores for some behaviors associated with stress (RAS, ESC) and negative odor valence (3’Z1) decreased, the scores for behavioral parameters associated with positive odor valence (e.g., dH/C, 1’dOI, 3’dOI, 3’Z3) showed marked increases.

Next, we performed a hierarchical clustering analysis on the behavioral score matrix to assess how all the odorants at the different concentrations cluster in a multidimensional space. This analysis separated all the odorants and concentrations tested into two groups (Fig. 5C). The first group is composed mostly (7/8) of odorants presented at the highest concentration (85 mM). Of those eight odorants, three odorants elicit aversion, and five are neutral. In contrast, the second group includes H2O, eight neutral, and 20 approached odorants. In that group, five odorants (TMA, DMP, PUT, TMT, and VAN) produce behaviors which, while diverse across odorants, were quite similar across the two lowest concentrations of the same odorant. However, all other odorants are not tightly clustered across concentrations of themselves, suggesting that the behavioral profile is a function of concentration. A PCA on these data further supported these results (Fig. S5D).

Finally, to compare the effects of odorant identity vs. odorant concentration, we calculated the Euclidean distances between all possible odorant-concentration pairs. We found that the average distances for different odorant pairs of equal concentrations (i.e., inter-odorant), and for different concentration pairs of the same odorant (i.e., inter-concentration) are not significantly different (Fig. 5D). However, the average inter-odorant distances were significantly higher for odorant pairs at the highest concentration (85 mM) compared to odorant pairs at the two lower concentrations (850 μM and 8.5 μM). Together, these data suggest that odorant identity and concentration have roles of a similar magnitude in determining mouse olfactory behavior, and that higher odorant concentrations increase the diversity of odor-guided behaviors across odorants.

## DISCUSSION

Over the last 30 years, some studies have contributed to the quantification, characterization, and even the prediction of the human olfactory percept. This was made possible by applying a combination of modern tools for the physicochemical characterization of odorants, statistical methodologies, and machine learning algorithms to large human olfactory psychophysical datasets (6–11). In comparison, the mouse olfactory percept is still poorly understood, mainly due to the lack of a large dataset for murine odor-guided behaviors. Here, we conducted a large-scale study of the effects of odorant exposure on mouse behavior and generated the most comprehensive mouse olfactory ethological atlas to date. By scoring 18 different behaviors in 525 mice across 98 odorant conditions, we generated 9765 data points encompassing 1764 odorant-behavior interactions. This atlas uncovered many relationships between odorants and odor-guided behaviors, and allowed us to: i) characterize mouse olfactory behavior for 73 odorants in unprecedented detail, ii) understand how mouse behavior changes across concentration, iii) use the physicochemical properties of odorants to predict odor-guided behaviors in mouse, and iv) describe a weak relationship between mouse and human olfactory percepts.

In humans, odor character is quantified by measuring behavioral responses to odors, using methods such as free labeling, odor profiling, or pairwise similarity (39). These methods show that, although there is variation across individuals, humans have a shared and reproducible olfactory percept for a given odor. Similarly, we find that mouse olfactory perception can be quantified by measuring behavioral responses, and that these behaviors demonstrate a shared and reproducible olfactory percept across individuals. Many odorants were clearly distinguishable even in a simple low-dimensional space built from these behaviors and, more formally, many individual odorants could be discriminated from each other or from the ensemble, even in a novel mouse. Odor behavior in novel mice could even be partly reconstructed using a low-dimensional space built from the remainder of the dataset. Furthermore, a single dimension showed a modest correlation between mouse behavior and human perception, which was absent in any higher dimensions. Lastly, some mouse behaviors were at least partly predictable from chemical structure alone. Together, these data indicate that there is indeed a rich, canonical set of odor-evoked behaviors in mouse, and this analysis begins to give us some insight into how mice perceive odor character.

The olfactory system employs a combinatorial strategy to maximize its detection and discrimination power of different odorants across various concentrations (30, 31). Depending on the species, this strategy involves hundreds to thousands of OSN/OR subtypes present in the olfactory mucosa at different abundances (3), responding to potentially millions of odorants in the environment. Given these numbers, only a combinatorial olfactory behavioral code would allow the diverse content of the olfactory world and the physiological response to it to be accurately reflected in the actions of an animal. In other words, each odorant should be able to trigger more than one behavior, each behavior should be triggered by many odorants, and different odorants would trigger different sets and magnitudes of behaviors. Indeed, three key results in this study reflect the combinatorial nature of the olfactory system. First, the vast majority of the odorants and concentrations induced unique behavioral codes. Second, we noted that distinct combinations of physicochemical properties predicted different behaviors. Third, we found that the number of significant behaviors decreases with descending concentrations of odorants, consistent with the fact that lower odorant concentrations activate lower numbers of mouse ORs and smaller numbers of glomeruli in the olfactory bulb (30, 32, 40), theoretically allowing a smaller number of combinatorial activity patterns in downstream brain areas.

The results above raise the hypothesis that in addition to being combinatorial, the mouse olfactory perceptual space would also be highly multidimensional. However, our data were consistent with a low-dimensional space for behavior. While this does not prove that the perceptual space is also low-dimensional, it does not provide any evidence against it either. Odor valence, which in humans is the principal perceptual dimension by which odors are categorized (7), emerged as the most important behavioral dimension across our analyses, as it segregated our data into two major groups associated with behaviors indicating either a positive or negative valence. Thus, we conclude that odor valence is the primary axis of olfactory perception in mouse, similar to what has been reported in human (7, 12, 26). Together, these results provide additional lines of evidence to the hypothesis that the dimensionality at the behavioral/perceptual levels is indeed lower than it first appears (6).

The impact of concentration on mouse olfactory preferences has been analyzed in multiple studies (14, 25, 37, 41, 42). In this study, we extended these analyses to 18 behaviors for 12 different odorants at three different concentrations, yielding several novel findings, and making this the most comprehensive study so far on the effects of odorant concentration on mouse olfactory behavior. We found that different concentrations elicit different behavioral combinatorial codes, and that the highest concentration elicits the highest number of significant changes in behaviors. While we cannot exclude the fact that the lower concentration is too weak to be detected by the mice, two observations argue against that – first, we observe significant changes behavioral changes for all odorants but PEA at the lowest concentration, and second, PEA activates TAAR-expressing OSNs at concentrations several orders of magnitude lower than the lowest one used in this study (43).

Additionally, we demonstrated that odor character is not an intrinsic property of a molecule, but rather changes across concentration. These findings are consistent with results from human studies, which showed that higher concentrations elicit the most apparent differences in odor quality between odorants and that odorants at lower concentrations show higher degrees of similarity between their odor qualities (36). Together, the data presented here support a model where the olfactory percepts of mouse and human share many common properties, despite displaying different olfactory preferences.

Multiple studies have linked the molecular features of odorants to psychophysical and behavioral measures of odor valence in human and mouse, respectively (7, 17, 34). Moreover, a recent international crowdsourcing competition was able, for the first time, to accurately predict the human olfactory percept of many odorants (8). To successfully achieve that goal, the researchers combined machine learning algorithms with a large human psychophysical dataset collected from 49 individuals profiling 476 odorants. In the present study, we employed a similar strategy and found that some mouse odor-guided behaviors can also be partly predicted by the physicochemical properties of the odorants. The most predictable of these were the valence-related behaviors (i.e., 1’dOI and 3’dOI), which were also among the ones displaying the lowest interindividual variability across all odorants.

The observation that specific physicochemical properties of odorants can predict some behavioral outputs in mouse opens new possibilities for studying the functional organization of the mouse olfactory system. These results raise the hypothesis that other subsets of physicochemical properties predictive of odor-guided behaviors could potentially be linked to the spatial organization of ORs/OSN subtypes in the mouse nose. Future large-scale experiments focused on connecting the zonal expression patterns for all mouse ORs to the physicochemical descriptors of their respective agonists will be critical to test this hypothesis.

In conclusion, our study provides a foundational quantitative database of odor-guided behaviors in the mouse that can be exploited in future studies to further deconstruct many aspects of the mouse olfactory percept, and facilitate future comparative studies of olfactory percepts among different species.

## MATERIAL AND METHODS

### Behavioral Scoring

We retrieved a video library from our previous study (25), in which we subjected adult male C57Bl/6J mice (aged 8-14 weeks, The Jackson Laboratory) to the olfactory preference test for a total duration of 3 minutes (experiments were approved by the Fred Hutchinson Cancer Research Center Institutional Animal Care and Use Committee). For this study we analyzed 410 videos from mice exposed to an odorless control (i.e., water, or H2O) or one of the 73 odorants at a single concentration (at 85 mM), and 123 additional videos from mice exposed to two other descending concentrations (850 μM and 8.5 μM) for a subset of 12 odorants.

In addition to the cumulative duration of olfactory investigation (3’dOI) reported in (25) for the 3 minutes-long assay, we scored 17 new behavioral parameters indicative of either valence, stress, and exploration. The videos were either randomized and scored blind (to the odorant) using the following criteria:

- Olfactory investigation (OI) parameters: 1’dOI, 1’fOI, and 3’fOI represent the cumulative duration (d) or frequency (f) of olfactory investigation during the 1st minute (1’) or the full 3 minutes (3’) of the assay. We considered OI only if the nose of the mouse was overlapping, or in very close proximity (~0.5 cm) of the stimulus. For these parameters, the videos were randomized and scored by the experimenter.
- Zone (Z) parameters: 1’Z1, 1’Z3, 3’Z1, and 3’Z3 represent the cumulative time mice spent in either Zone 1 (Z1) or Zone 3 (Z3) of the cage during the 1st minute (1’) or the full 3 minutes (3’) of the assay. Here, the test cage was divided into three equal-sized zones, with Z1 representing the zone furthest away from the odor stimulus and Z3 the zone containing the stimulus. Time spent inside each zone (head of the animal had to be within the zone) was scored using the Ethovision XT software (version 11, Noldus Information Technology), and videos in which the mouse transported the piece of filter paper outside Z3 were not included. For these parameters, the videos were scored by a non-experimenter.
- Handling (H) or catching (C) parameters: dH/C and fH/C represent the cumulative duration (d) or frequency (f) where mice handled and/or caught the stimulus with its front paws during the 3 minutes of the assay. For these parameters, the videos were scored by a non-experimenter.
- Risk assessment (RAS): this parameter represents the number of episodes the mouse displays the flat-back/stretch-attend response, followed by a sniff in the direction of the stimulus, during the 3 minutes of the assay. For this parameter, the videos were scored by a non-experimenter.
- Escape (ESC): this parameter represents the number of episodes the mouse displays a quick and contactless approach towards the stimulus, followed by an even faster withdrawal/darting to the opposite end of the cage, during the 3 minutes of the assay. For this parameter, the videos were scored by a non-experimenter.
- Digging (DIG) parameters: dDIG and fDIG represent the cumulative duration (d) or frequency (f) that the mouse digs into the bedding with the forelimbs, often kicking it away with the hindlimbs, during the 3 minutes of the assay. For these parameters, the videos were scored by a non-experimenter.
- Distance (DIT): this parameter represents the total length the mouse walked/ran through during the three minutes duration of the video. Distance traveled was scored using the Ethovision XT software (version 11, Noldus Information Technology), and video tracking done using the center-point of the mouse. For this parameter, the videos were scored by a non-experimenter.
- Rearing (REA): this parameter represents the number of episodes the mouse stands on its hindlegs (rearing) anywhere in the cage, including when rearing against the walls during the 3 minutes of the assay. For this parameter, the videos were scored by a non-experimenter.
- Self-grooming (SGR) parameters: dSGR and fSGR represent the cumulative duration (d) or frequency (f) where the mice are self-grooming, defined by when the mouse was licking its fur, grooming itself with the forepaws, or scratching any part of its body with any limb. For these parameters, the videos were scored by a non-experimenter.

Each mouse was exposed to only a single odorant, and thus the data for each odorant consisted of 5-10 mice each scored on 18 behaviors.

### Odorants

Odorants used in studies with mice were assigned to different structural classes (underlined), as shown below, followed by their 3-letter abbreviation in parentheses:

Alcohols: 2-phenylethanol (2PE); geraniol (GER); heptanol (HPO); hexanol (HXO); linalool (LIN); octanol (OCO); cis-3-hexenol (C3H). Aldehydes: benzaldehyde (BZL); citral (CIT); citronellal (CTN); heptanal (HPN); octanal (OCN); trans-2-hexenal (T2H). Amines: 2-methylbutylamine (2MB); 1-(2-aminoethyl)piperidine (AEP); aniline (ANI); 3-amino-s-triazole (AST); cadaverine (CAD); N,N-dimethylbutylamine (DMB); N,N-dimethylethylamine (DME); N,N-dimethyloctylamine (DMO); N,N-dimethylcyclohexylamine (DMC); heptylamine (HEP); hexylamine (HXA); isoamylamine (IAA); 2-methyl-1-pyrroline (M1P); N-methylpiperidine (NMP); octylamine (OCT); 2-phenylethylamine (PEA); trimethylamine (TMA); putrescine (PUT); pyrrolidine (PYR); spermidine (SPD); spermine (SPN); o-toluidine (TOL). Azines: 2,5-dimethylpirazine (DMP); 2-ethyl-3,5(6)-dimethylpirazine (EDM); indole (IND); quinoline (QUI); skatole (SKA). Camphors: (+/−)-camphor (CAM); (−)-fenchone (-FCH); (+)-fenchone (+FCH); eucalyptol (EUC). Carboxylic Acids: 2-methylbutyric acid (MBA); octanoic acid (OCA); propionic acid (PPA). Esters: amyl acetate (AAC); ethyl butyrate (EBT). Ketones: 2-heptanone (2HO); alpha-ionone (ION). Musks: ambrettolide (AMB); civettone (CIV); muscone (MUS). Terpenes: beta-farnesene (BFA); (−)-carvone (-CVN); (+)-carvone (+CVN); farnesene mixed isomers (alpha+beta) (FAR); (−)-limonene (-LIM); (+)-limonene (+LIM); (+)-menthol (+MEN); (−)-menthone (-MNT); (+)-menthone (+MNT); alpha-pinene (PIN); rose oxide (ROX). Thiazoles: 2-isobutylthiazole (IBT); 2-isopropyl-4-5-dihydrothiazole (IPT); 2,5-dihydro-2,4,5-trimethylthiazoline (TMT). Thiols: 2-propylthietane (2PT); hexanethiol (HXT); octanethiol (OTT). Vanillin-like compounds: eugenol (EUG); vanillin (VAN).

### Machine Learning

We calculated 4,870 physicochemical descriptors for 74 molecules using Dragon 6.0 (Talete) software. We preprocessed the descriptors using the caret package and R Statistical Software (version 3.5.1). We first removed descriptors with near-zero variance, missing data, or that were highly correlated with another descriptor (>95%) leaving a final total of 662 features.

All available data were split 75%/25% between training and test sets. Support vector machine models were optimized for each of the 18 outcome variables using three different seeds and five-fold cross-validation with two repetitions. The mean predictive accuracy (R squared) across all runs is shown in (Data File S2 and Fig. 4E). For the best-performing outcome variable (3’dOI), we further examined the important Dragon descriptors, and plotted the top 20 (Fig. S4A). After ranking all of the variables on their importance in the model, we sequentially added 50 variables at a time until reaching the full model of 662 features (Fig. S4B). We also examined how the number of training examples available would influence training and test performance. We tested multiple subsets of 330 training examples against a hold-out set of 80 test examples to determine the training set size where the improvement in model performance plateaus (44). The learning curves for the Linear Support Vector Machine model are shown in Fig. S4C.

### DREAM predictions

The DREAM model developed in Keller et al, 2017 was applied to water and each of the 73 odorants used in the current study to generate predictions for each of 21 perceptual descriptors. Briefly, an isomeric, canonical SMILES string was generated for each odorant using rdkit (Python) and used to generate an rdkit mol object in which 3-dimensional coordinates of each atom position were estimated. Dragon 6.0 was used to compute features from these 3-dimensional structures. The DREAM model was re-trained from scratch on only 448 out of the 476 original molecules to avoid overlap with the molecules from the current study. It was then used to predict 21 perceptual descriptors from the 74 odorants used in the current study.

## Data Analysis

Some statistical analyses were done using Python 3, Pandas, and Scikit-Learn, GraphPad Prism (version 8.0.0), and PAlaeontological STatistics (version 4.06, http://folk.uio.no/ohammer/past/). Hierarchical clustering analysis was performed using Euclidean distances with Ward’s method, after data values were standardized. For principal component analysis, the data matrix was standardized and correlation matrixes used to compute the eigenvalues and eigenvectors. P-values for odorant/behavior interactions were computed with a two-stage Benjamini, Krieger and Yekutieli (BKY) multiple comparisons correction (45).

## ACKNOWLEDGMENTS

We would like to thank Dr. Darren W. Logan and the members of the Saraiva Lab for the constructive feedback. This work was supported by Sidra Medicine, and the National Institute of Health (U19NS112953).

## AUTHOR CONTRIBUTIONS

D.M. participated in the design of the project, analyzed data, and wrote the initial version of the manuscript. M.M., C.J.A., R.H., A.S., S.D., D.A., and S.S. analyzed data. J.D.M. analyzed data, and helped write the manuscript. R.C.G. designed the data analysis methodology, analyzed data, and wrote the final version of the manuscript. L.R.S. analyzed data, conceived and supervised the project, and wrote the final version of the manuscript.

## SUPPLEMENTARY MATERIAL

**Figure S1.**
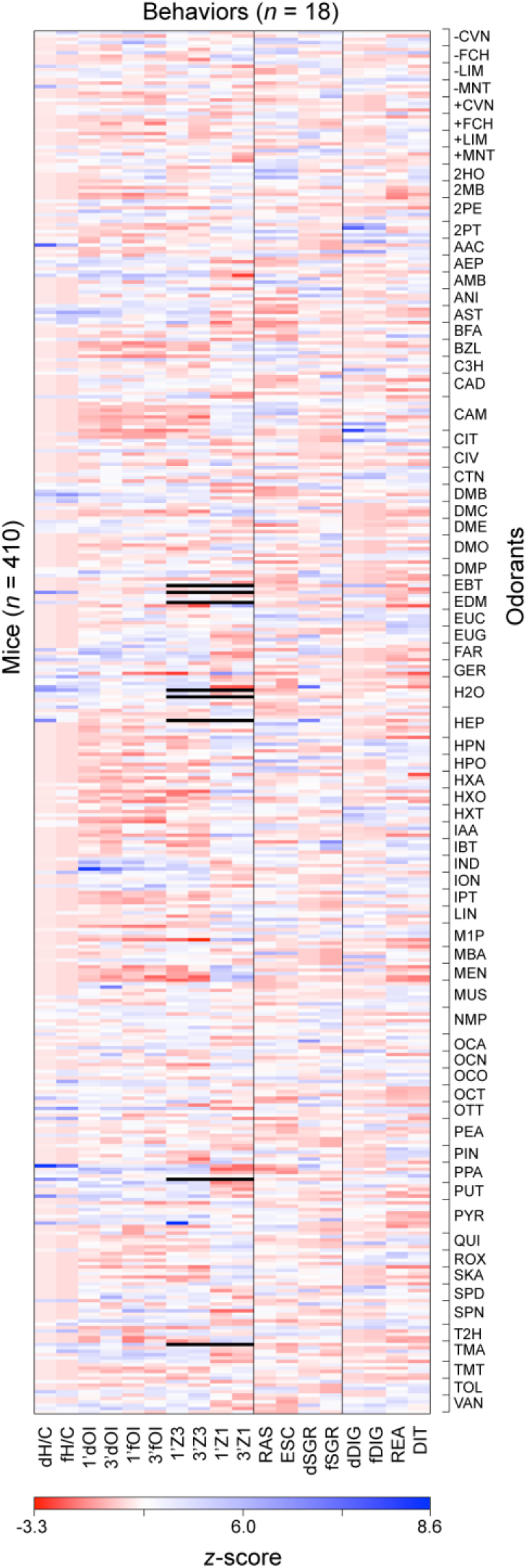
A mouse olfactory-ethological atlas. (**A**) Heatmap summarizing the 7380-individual z-scores representing of 18 mouse behavioral responses to 73 odorants (*n* = 410 mice).

**Figure S2.**
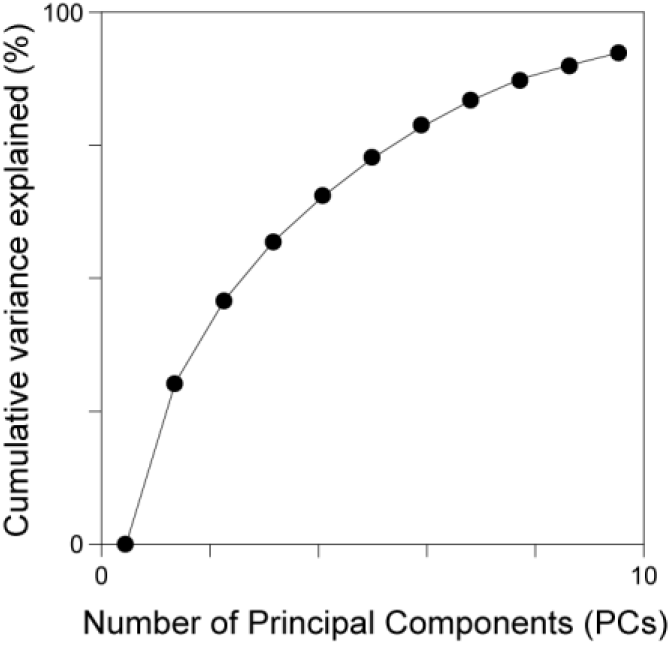
The primary axis of olfactory perception in the mouse. Cumulative variance explained by the principal components (PCs).

**Figure S3.**
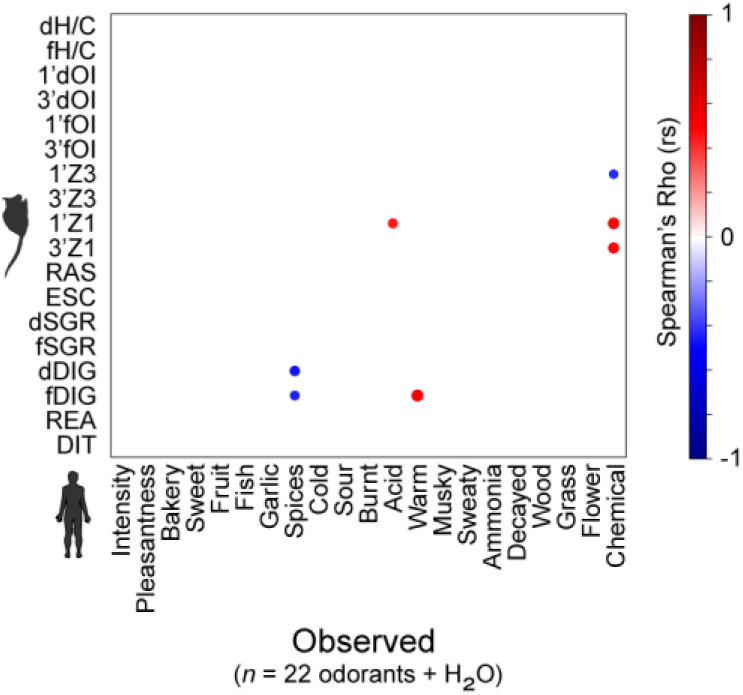
Comparing the mouse and human olfactory percepts. Correlogram between the 18 mouse behavioral parameters and 21 human ratings for the 23 odorants overlapping between this study and (*9*). Only significant (P < 0.05) correlations are plotted, and the circle size and color indicate the magnitude and direction of the correlation (Spearman rho, rs). Blank cells correspond to non-significant correlations.

**Figure S4.**
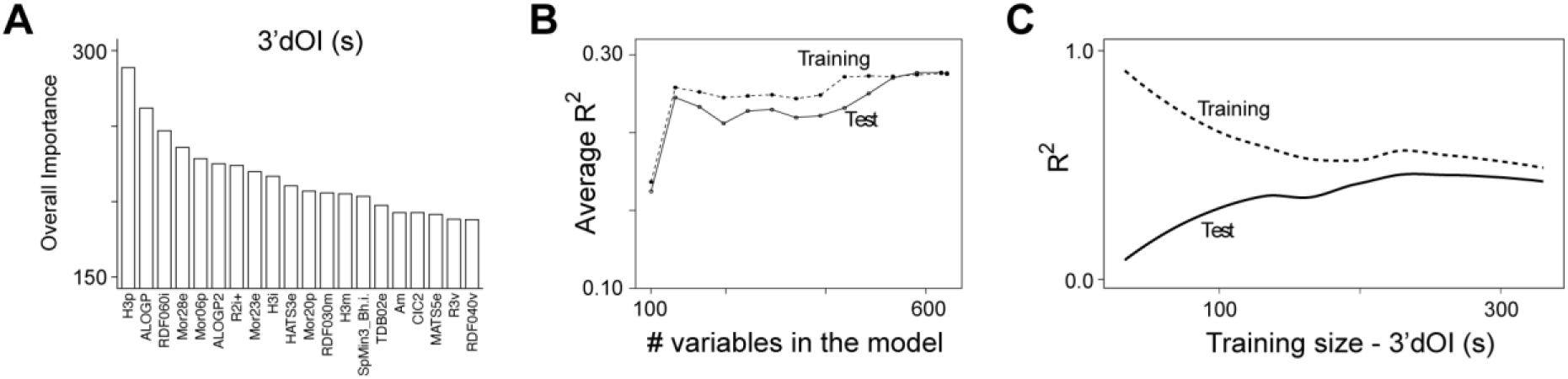
Predicting mouse olfactory-driven behaviors. (**A**) Histogram showing twenty physicochemical descriptors with highest importance rating for 3’dOI. (**B, C**) Model performance as a function of the number of variables in the model (B) and number of odors in a resampled training set (C).

**Figure S5.**
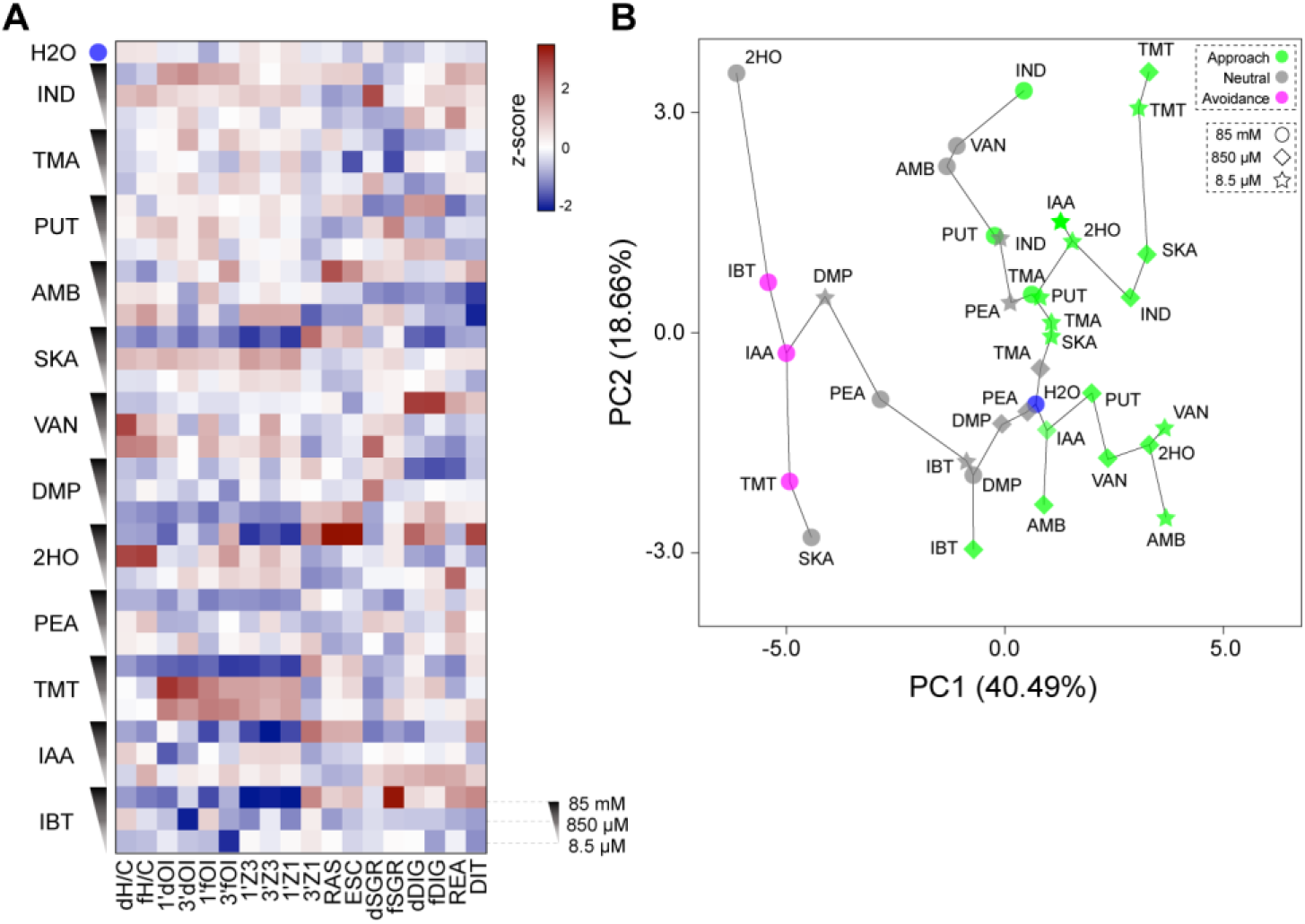
The effect of odorant dosage on behavior. (**A**) Heatmap indicating the z-score for each of 12 odorants (relative to all other odorants) for the 18 behavioral parameters measured after exposure to those odorants at three concentrations (85mM, 850μM, and 8.5μM). (**B**) Principal component analysis of the behavioral profiles for the 12 odorants tested at the three different concentrations. H2O is shown in blue, aversive odorants in magenta, neutral in grey, and approached in green. The line connecting the circles indicates the minimum spanning tree.

**Data file S1** (Microsoft Excel format). Raw behavioral scores for the 410 mice exposed to odorants; P-values for the odorant vs. H2O comparison for each of the behaviors (one-way ANOVA, BKY multiple comparisons correction): ns, non-significant; *P < 0.05; ***P < 0.001. Behaviors showing significant increases and decreases are highlighted in green and magenta, respectively. Grey color indicates no significant statistical difference.

**Data file S2** (Microsoft Excel format). Linear regression and correlation stats for 3’dOI vs. the number of carbons; Spearman correlation coefficient (rs) matrix of the 18 behavioral parameters vs. 1536 P-C descriptors; R^2^ matrix of the machine learning training vs. test.

**Data file S3** (Microsoft Excel format). Raw behavioral scores for 36 odor conditions and H_2_O across 18 behavioral parameters; P-values of 36 odor conditions across 18 behavioral parameters vs. H_2_O control (one-way ANOVA, BKY multiple comparisons correction); Euclidean distances between 36 odor conditions and H_2_O, across 18 behavioral parameters.

